# Single-Cell Sequencing Identifies the Crucial Role of Mitochondrial Fission-Fusion Balance in Cardiac Hypertrophy Progression

**DOI:** 10.1101/2025.02.13.638057

**Authors:** Tao He, Jianmei Sha, Yuxin Hu, Caihong Shao, Yi Zhou, Lu Chen, Jianhua Yao, Junli Gao

## Abstract

**Background:** The heart undergoes growth in response to both pathological and physiological stimuli. Pathological hypertrophy often leads to cardiomyocyte loss and heart failure (HF), whereas physiological hypertrophy paradoxically protects the heart and enhances cardiomyogenesis. The molecular mechanisms that distinguish these two forms of hypertrophy remain unclear.

**Methods:** In this study, we utilized single-cell transcriptomics from transverse aortic constriction (TAC) models at 2, 5, 8, and 11 weeks (GSE120064), along with bulk RNA sequencing from mice subjected to 12 months of exercise-induced physiological hypertrophy and cardiomyogenesis (CRA007207), to investigate the molecular differences between pathological and physiological hypertrophy.

**Results:** Our results reveal the following. Mitochondrial-related pathways are the primary drivers of the pathological changes that occur following TAC. The mitochondrial fission and fusion pathways exhibited increased activity at 2 weeks but decreased activity at 5, 8, and 11 weeks post TAC. The expression pattern of exercise-induced physiological hypertrophy was similar to that of 2-week TAC-induced changes, indicating that the early stage of TAC represents an adaptive physiological response or physiological hypertrophy. Notably, during HF, the fission genes Fis1 and Dnm1l increase, in contrast to the expected decrease in fusion genes. These findings were experimentally validated, indicating that the mitochondrial fission genes Fis1 and Dnm1l are key promoters of HF.

**Conclusions:** Our data indicate that the balance between mitochondrial fission and fusion plays a critical role in the transition from physiological to pathological hypertrophy. The fission-related genes Fis1 and Dnm1l have emerged as key drivers of pathological hypertrophy and heart failure. These findings suggest that targeting fission genes, particularly Fis1 and Dnm1l, may represent promising therapeutic strategies for managing heart failure.

## Introduction

Cardiovascular disease (CVD) remains one of the leading causes of mortality worldwide, with heart failure representing a significant and growing burden on public health^1^. The prevalence of HF increases with age, and despite advances in medical care, the prognosis remains poor, with a 5-year mortality rate reaching 50%^2^.

Physical activity is associated with a reduced risk of HF. In animal models, exercise training paradoxically alleviates dysfunction and fibrosis typically induced by pathological stimuli. The cardiac benefits of physical exercise are widely recognized. Researchers are investigating the changes that occur in the heart after exercise and believe that gene variations induced by exercise could serve as therapeutic targets for cardiac diseases, including HF.

Exercise-induced cardiac hypertrophy is a physiological response, whereas pathological hypertrophy, despite sharing a similar phenotype, leads to HF. Some researchers propose that the key difference between the two lies in the nature of the stimulus: exercise is intermittent, while disease-induced stress is constant. Initially, compensatory hypertrophy helps maintain cardiac output under pressure overload, but prolonged stress can turn this adaptive response into a maladaptive one, worsening heart function and contributing to HF. Understanding the molecular mechanisms that drive this transition is crucial for identifying potential therapeutic targets to prevent or delay HF progression.

Research has identified distinct gene expression profiles between pathological and physiological hypertrophy, with early divergence seen in markers such as ANP, BNP^4^, PGC1α^5^, C/EBPβ^3^, and MYH6/7^6^, as well as noncoding RNAs (miR-222^7^, circDdx60^8^ and lncExACT1^9^). These markers are frequently used to differentiate between physiological and pathological hypertrophy.

Mitochondrial dynamics, encompassing the processes of fission and fusion, are critical for maintaining heart health^10^. As the powerhouses of cells, mitochondria preserve their functionality through continuous cycles of biogenesis, fission, fusion, and degradation^11^. Although relatively immobile in the adult heart, the morphological changes they undergo, driven by mitochondrial morphology factors, are vital for cellular processes such as energy production, organelle integrity, and stress responses^12^. Mitochondrial fusion proteins, particularly Mfn1/2 and Opa1, not only facilitate fusion but also play roles in endoplasmic reticulum tethering, mitophagy, cristae remodeling, and the regulation of apoptosis. Conversely, mitochondrial fission, governed by proteins such as Dnm1l, Fis1, Mff, and MiD49/51, is crucial for the removal of damaged mitochondria via mitophagy and for proper cell division^13^. Dysregulation of mitochondrial dynamics in the cardiac system is the key to HF^14^.

In this study, by combining scRNA-seq data from TAC mice at five different time points with bulk RNA-seq data from exercised mouse hearts, we found that the balance between mitochondrial fusion and fission is tightly controlled in cardiomyocytes during both pathological and physiological hypertrophy^15,16^. Our analysis revealed that mitochondria-related pathways are key drivers of gene expression changes over time following TAC. Specifically, the mitochondrial fission and fusion pathways were upregulated at 2 weeks post TAC, but this activity decreased at 5, 8, and 11 weeks. Weighted gene coexpression network analysis (WGCNA) further revealed that exercise-induced physiological hypertrophy is similar to the cardiac hypertrophy induced by 2 weeks of TAC, suggesting that 2 weeks of TAC is essentially compensatory hypertrophy or physiological hypertrophy. Despite these shared trends in mitochondrial dynamics, fission-related genes behave differently between the two conditions. Notably, in HFs, the fission genes Fis1 and Dnm1l are upregulated, whereas the fusion genes are downregulated as expected. The experimental validations suggest that the mitochondrial fission genes Fis1 and Dnm1l play critical roles in the transition from physiological to pathological hypertrophy in cardiomyocytes. Therefore, promoting mitochondrial fusion while selectively inhibiting fission, particularly by targeting Fis1 and Dnm1l, may represent a promising therapeutic strategy for treating cardiac diseases.

## Methods

### Animal studies

For this study, 8-week-old male C57BL/6J mice were used. All animal procedures were performed in accordance with protocols approved by the Institutional Animal Care and Use Committee (IACUC) of Shanghai University, China.

### Single cell RNA sequencing (scRNA seq) analysis

Single-cell RNA sequencing data from heart tissues of transverse aortic constriction model mice were obtained from the Gene Expression Omnibus (GEO) database (GSE120064). Cells expressing fewer than 200 or more than 10,000 genes were excluded to remove noncellular debris and potential cell aggregates. The data were log-normalized, and highly variable genes were identified via the FindVariableFeatures function. Subsequent analyses, including data scaling and principal component analysis (PCA), were conducted via the ScaleData and RunPCA functions, respectively. Cell clustering was performed with the FindNeighbors and FindClusters functions, and uniform manifold approximation and projection (UMAP) was used for cluster visualization, implemented in the Seurat R package as per the official vignettes (https://satijalab.org/seurat/articles/get_started.html). Cluster annotation was performed on the basis of known cell markers.

### Mitochondrial fission and mitochondrial fusion scores

To assess the activity of the fusion and fission gene sets in individual cells, the AddModuleScore function in Seurat was applied. The gene sets for mitochondrial fission and fusion were derived from the Gene Ontology (GO) terms ‘mitochondrial fission’ (GO:0007006) and ‘mitochondrial fusion’ (GO:0000266), respectively (Table 1).

**Table 1.**
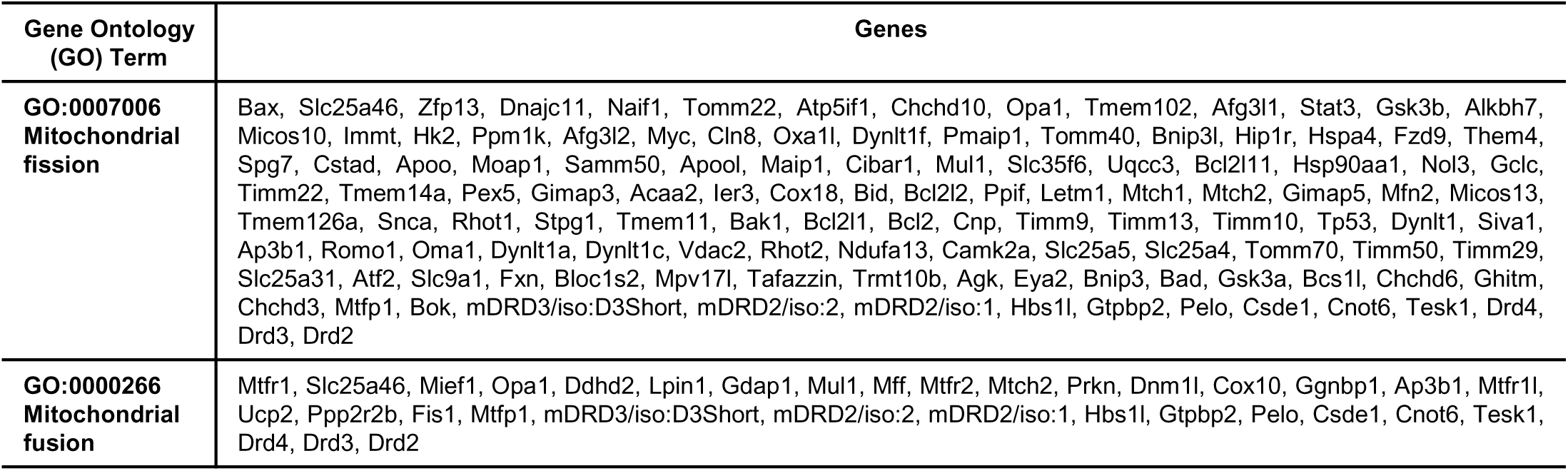

| Gene Ontology (GO) Term | Genes |
| --- | --- |
| GO:0007006 Mitochondrial fission | Bax, Slc25a46, Zfp13, Dnajc11, Naif1, Tomm22, Atp5if1, Chchd10, Opa1, Tmem102, Afg3l1, Stat3, Gsk3b, Alkbh7, Micos10, Immt, Hk2, Ppm1k, Afg3l2, Myc, Cln8, Oxa1l, Dynlt1f, Pmaip1, Tomm40, Bnip3l, Hip1r, Hspa4, Fzd9, Them4, Spg7, Cstad, Apoo, Moap1, Samm50, Apool, Maip1, Cibar1, Mul1, Slc35f6, Uqcc3, Bcl2l11, Hsp90aa1, Nol3, Gclc, Timm22, Tmem14a, Pex5, Gimap3, Acaa2, Ier3, Cox18, Bid, Bcl2l2, Ppif, Letm1, Mtch1, Mtch2, Gimap5, Mfn2, Micos13, Tmem126a, Snca, Rhot1, Stpg1, Tmem11, Bak1, Bcl2l1, Bcl2, Cnp, Timm9, Timm13, Timm10, Tp53, Dynlt1, Siva1, Ap3b1, Romo1, Oma1, Dynlt1a, Dynlt1c, Vdac2, Rhot2, Ndufa13, Camk2a, Slc25a5, Slc25a4, Tomm70, Timm50, Timm29, Slc25a31, Atf2, Slc9a1, Fxn, Bloc1s2, Mpv17l, Tafazzin, Trmt10b, Agk, Eya2, Bnip3, Bad, Gsk3a, Bcs1l, Chchd6, Ghitm, Chchd3, Mtfp1, Bok, mDRD3/iso:D3Short, mDRD2/iso:2, mDRD2/iso:1, Hbs1l, Gtpbp2, Pelo, Csde1, Cnot6, Tesk1, Drd4, Drd3, Drd2 |
| GO:0000266 Mitochondrial fusion | Mtfr1, Slc25a46, Mief1, Opa1, Ddhd2, Lpin1, Gdap1, Mul1, Mff, Mtfr2, Mtch2, Prkn, Dnm1l, Cox10, Ggnbp1, Ap3b1, Mtfr1l, Ucp2, Ppp2r2b, Fis1, Mtfp1, mDRD3/iso:D3Short, mDRD2/iso:2, mDRD2/iso:1, Hbs1l, Gtpbp2, Pelo, Csde1, Cnot6, Tesk1, Drd4, Drd3, Drd2 |

### Enrichment analysis

For pathway enrichment analysis, cluster-specific markers or differentially expressed genes identified via the FindMarkers function in Seurat were subjected to gene set variation analysis (GSVA) or GO enrichment analysis in RStudio. The R package clusterProfiler was used for enrichment analysis, with a q value cutoff of 0.05. The results were visualized as bar plots generated with ggplot2, and volcano plots, which were used to compare differential gene expression between groups, were created via ggplot2 in R.

### WGCNA Analysis

Bulk RNA sequencing data from heart tissues of exercise model mice were obtained from the GEO database (CRA007207). Gene coexpression analysis was performed via WGCNA with the WGCNA package in R. First, an adjacency matrix was generated from the gene expression matrix, and the coexpression relationships between gene pairs were calculated via Pearson correlation coefficients. Soft thresholding power (β) was applied to strengthen the adjacency matrix, with the β value selected on the basis of the scale-free topology criterion for each dataset.

Next, a topological overlap matrix (TOM) was constructed from the strengthened adjacency matrix to assess gene pairwise relationships, and genes with similar expression profiles were grouped into gene modules via average linkage hierarchical clustering on the basis of TOM dissimilarity (1-TOM). Gene modules were identified through dynamic tree cutting, and modules with at least 75% similarity were merged. The representative gene of each module, known as the module eigengene (ME), was calculated, and the correlation between the ME and each gene in the module was defined as the module membership (MM).

To assess differential gene expression between exercise (Run) and control heart tissues, p values were computed via a t test, and gene significance (GS) was determined via log10 transformation of the p value. The average GS of genes in a module was defined as the module significance (MS), which reflects the association between the module and the exercise condition.

### Expression Validation of Key Genes via Quantitative PCR

To validate the expression of key genes identified through bioinformatics analysis between myocytes and normal tissue, transcriptomes from 4 exercise (Run) mice and 4 sedentary (Sed) mice were analyzed. Total RNA was extracted from heart tissues via TRIzol reagent (Invitrogen, USA), followed by reverse transcription into cDNA via the PrimeScript RT Master Mix (Takara, Japan). Quantitative PCR (qPCR) was performed via a SYBR Premix Ex Taq II kit (Takara, Japan). Gene expression levels were normalized to that of GAPDH, which was used as the internal control. The sequences of primers used for qPCR were as follows.

Mfn1

forward 5′ - GAGAAACATTTTTTCCATAAGGTG - 3′

reverse 5′ - CTTCTGGCATCCCCTGAGCTTTATG - 3′

Mfn2

forward 5′ - GTGGAAGGCAGTGGGCTGGAGACTC - 3′

reverse 5′ - CAGTTTGGCTCTGCTCTGAAGTG - 3′

Opa1

forward 5′ - CGACTAGAGAAAAACGTTAAAGAG - 3′

reverse 5′ - CACGTACATGACTTTTACCCTGTG - 3′

Dnm1l

forward 5′ - GTGGGATTGGAGACGGTGGTCAG - 3′

reverse 5′ - GCTTCTTTTCTTCGTTGGGCCATG - 3′

Fis1

forward 5′ - GAAAGGAAATTTCAGTCTGAGCAG - 3′

reverse 5′ - CATGGCCTTATCAATCAGGCGTTC - 3′

Gapdh

forward 5′ - CATCACTGCCACCCAGAAGACTG - 3′

reverse 5′ - ATGCCAGTGAGCTTCCCGTTCAG - 3′

### Statistical Analyses

Statistical analyses were performed via GraphPad Prism 8 unless otherwise stated. Unpaired, two-tailed Student’s t tests were used for comparisons between two groups at a single time point. For comparisons involving multiple groups, one-way ANOVA with Sidak’s or Tukey’s post hoc tests was applied, with multiplicity-adjusted p values reported for each comparison. Time-course data were analyzed via repeated-measures two-way ANOVA with Sidak’s post hoc test. Fisher’s exact test was used for event rate comparisons. The data are presented as the means ± SEMs, with statistical significance set at p < 0.05. The number of animals (n) represents the number of biological replicates used in each experiment. Sample sizes were determined on the basis of empirical data and expected data completeness to ensure sufficient power to detect differences, if present.

## RESULTS

### Mitochondria-Related Modules are Differentially Expressed in the TAC Mouse Model

To characterize the major cell types involved in cardiac remodeling following pressure overload, we performed clustering analysis and identified six key cell populations, including CMs, endothelial cells (ECs), fibroblasts (FBs), macrophages (MPs), T cells, and granulocytes (GNs), on the basis of distinct molecular markers (Fig. 1A, B).

**Figure 1.**
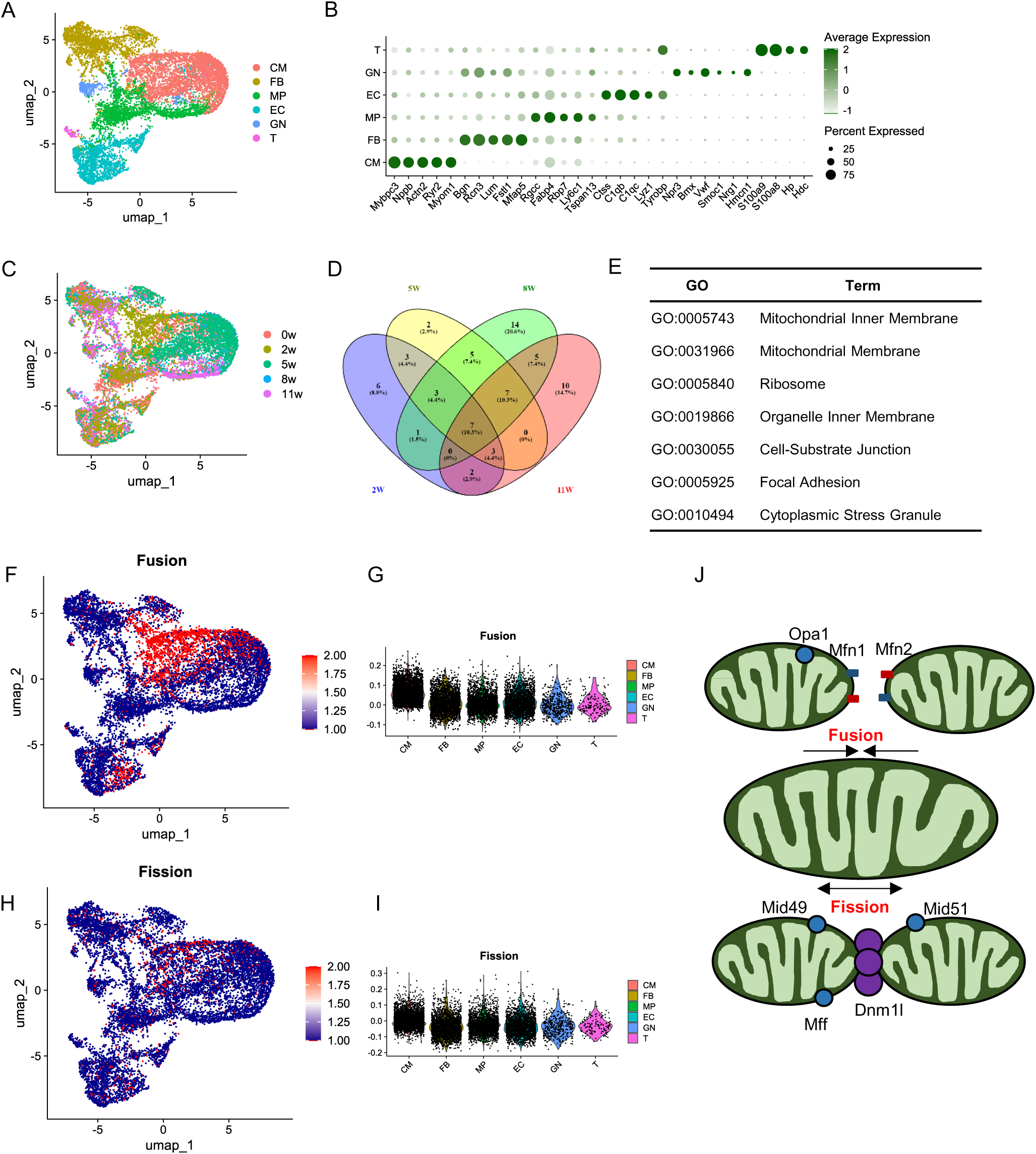
Mitochondria-Related Modules are Differentially Expressed in the TAC Mouse Model. **A.** UMAP plot illustrating the transcriptomic landscape, consisting of six major cell types. The cells are colored according to their clusters, with the following cell types: CMs (cardiomyocytes), ECs (endothelial cells), FBs (fibroblasts), GNs (granulocytes), MPs (macrophages), and T cells (T cells). **B.** Dot plot displaying DEGs across each cell type. **C.** UMAP plot showing the distribution of samples across different time points. **D.** Venn diagram illustrating the enriched GO terms for DEGs at different time points. **E.** Seven overlapping GO terms enriched at the post-TAC time point. **F, H.** Mitochondrial fusion/fission-related gene expression at various stages of cardiac hypertrophy. **G, I.** Violin plot demonstrating that fusion/fission-related genes are expressed primarily in CMs. **J.** Schematic diagram depicting mitochondrial fusion and fission. Mitochondrial remodeling involves two processes, fission and fusion, which are mediated by specific proteins. During fission, proteins such as Dnm1l, Fis1, and Mff become activated and localize to mitochondria, leading to mitochondrial fragmentation. This process either creates a new mitochondrion or separates dysfunctional mitochondrial components. In contrast, mitochondrial fusion is driven by outer membrane fusion proteins (Mfn1 and Mfn2) and the inner membrane fusion protein Opa1. The tethering of outer membranes, followed by the fusion of inner membranes, results in the formation of elongated mitochondria.

To assess the molecular changes associated with the progression of cardiac hypertrophy, we systematically examined hearts from different anatomic regions of the TAC mouse model at representative time points (0, 2, 5, 8, and 11 weeks) post TAC surgery (Fig. 1C).

We then conducted GO enrichment analysis to explore the pathways activated at different stages of hypertrophy. Seven GO terms were consistently enriched across all stages, including mitochondrion inner membrane, mitochondrial membrane, ribosome, organelle inner membrane, cell-substrate junction, focal adhesion, and cytoplasmic stress granule (Figure 1E). Notably, two of these terms were directly related to mitochondria. Given the critical role of mitochondria in the highly oxidative myocardium, maintaining mitochondrial function is essential for optimal cardiac performance.

Mitochondrial dynamics—including fusion, fission, biogenesis, and mitophagy—are fundamental for regulating mitochondrial morphology, quality control, and abundance, processes that have recently been implicated in cardiovascular disease. Our analysis focused specifically on mitochondrial fission and fusion (Fig. 1J). The key genes involved in mitochondrial fusion are dynamin-related GTPases termed mitofusins (Mfn1 and Mfn2) and opticatrophy protein 1 (Opa1), whereas those associated with mitochondrial fission are mitochondrial fission 1 protein (Fis1) and dynamin-related protein 1 (Dnm1l).

To elucidate the role of mitochondrial fission and fusion in the development of cardiac hypertrophy, we evaluated the expression levels of fission- and fusion-related proteins across different stages of disease progression.

A “mitochondrial fission and fusion score” was derived from the expression levels of proteins associated with the GO terms for mitochondrial fission (GO:0007006) and fusion (GO:0000266) (Table 1).

Further analysis revealed that these fission- and fusion-related proteins were predominantly expressed in CMs, with only limited expression observed in ECs, FBs, MPs, T cells, and GNs (Fig. 1F, I). Both violin and ridge plots further confirmed the enrichment of these proteins in CMs (Fig. 1G, H, I, J).

In summary, our findings demonstrate that mitochondrial-related modules are differentially expressed in the TAC mouse model, with a specific focus on mitochondrial fission and fusion processes. These processes are primarily active in cardiomyocytes, underscoring their potential importance in the development of cardiac hypertrophy.

### Variation in Mitochondrial Fission and Fusion Module Expression from Cardiac Hypertrophy to Heart Failure

Five time points were selected to represent different stages of cardiac hypertrophy following TAC-induced cardiac hypertrophy: 0, 2, 5, 8, and 11 weeks. The 0-week time point represents normal cardiac function, whereas 2 weeks indicates hypertrophy with a preserved ejection fraction (EF). After 2 weeks, the animals were allowed to stand for physiological hypertrophy, which had a protective effect on the myocytes. At 5 and 8 weeks, hypertrophy with reduced EF was observed, indicating pathological hypertrophy. By 11 weeks, heart failure was evident (Fig. 2A).

**Figure 2.**
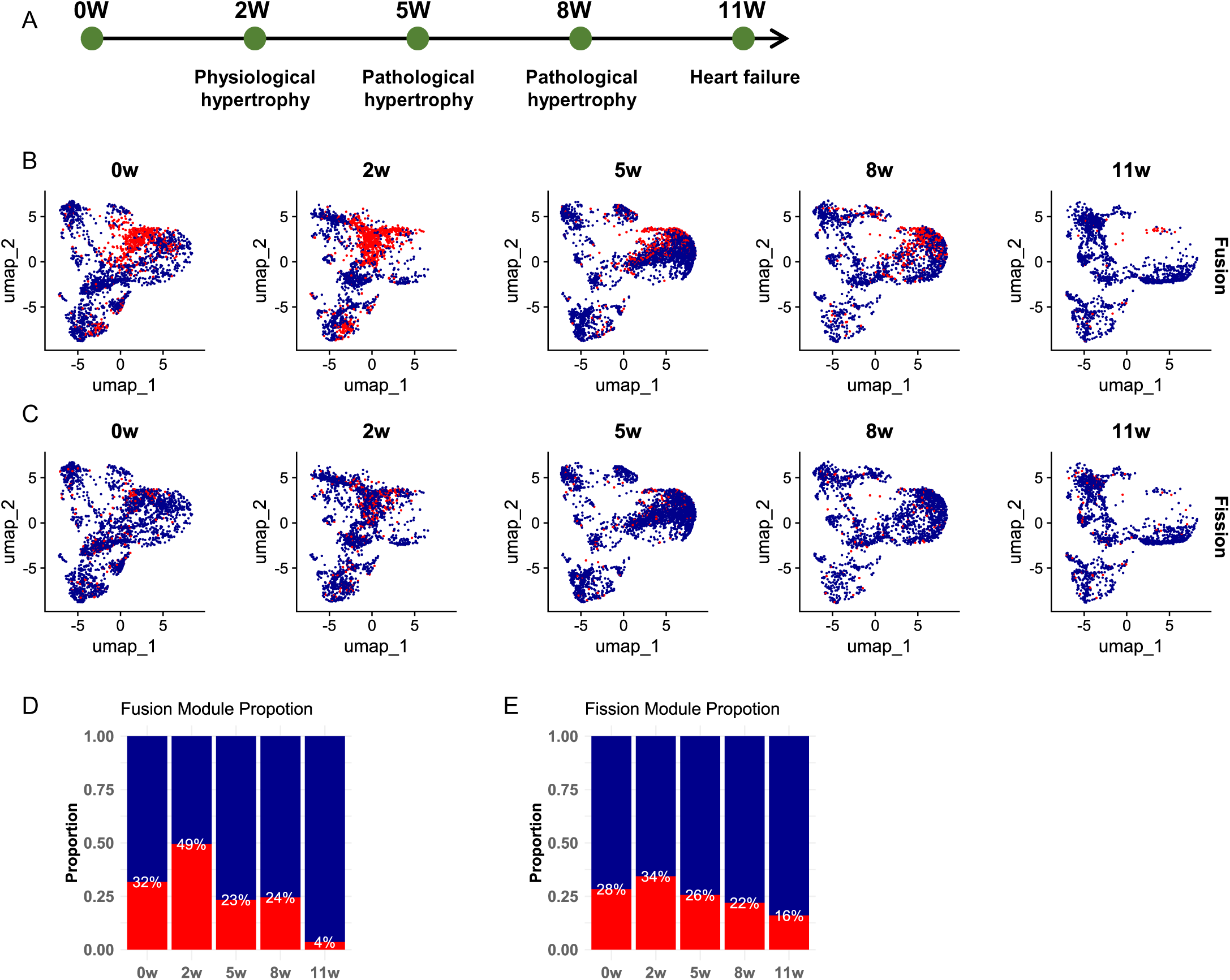
Mitochondria-Related Modules are Differentially Expressed in the TAC Mouse Model. **A.** Schematic representation of the study design, showing cardiac phenotypes at different time points. “2 W” represents physiological hypertrophy, whereas “5 W” and “8 W” correspond to stages of pathological hypertrophy. “11 W” marks the transition to heart failure. **B.** Expression patterns of fusion-related genes at different time points. **C.** Expression patterns of fission-related genes at different time points. **D.** Proportions and composition of genes involved in mitochondrial fission and fusion.

At 2 weeks, as the heart undergoes physiological hypertrophy, mitochondrial fusion increases from 32% to 49%. This finding suggests that the heart adapts to meet higher energy demands by enhancing mitochondrial dynamics to support its increased workload.

However, at 5 and 8 weeks, as pathological hypertrophy progresses with a decline in EF, energy demands decrease. Consequently, both mitochondrial fusion and fission decrease, reflecting the heart’s diminishing capacity for energy production.

By 11 weeks, during heart failure, the heart is severely impaired, and energy demands are minimal. Mitochondrial fusion decreased to 4%, whereas fission decreased to 16%, reflecting the reduced functional capacity of the heart at this stage (Fig. 2B, C, D, E).

In summary, during the early stages of TAC-induced hypertrophy, the heart compensates by increasing its energy production capacity to meet the body’s demands and has a protective function. However, as the condition progresses, cardiac function deteriorates, and energy requirements diminish, ultimately leading to irreversible damage and heart failure.

### Expression of Mitochondrial Fusion and Fission Genes During Cardiac Hypertrophy

As previously mentioned, the primary regulators of mitochondrial fusion in mammalian cells are Mfn1, Mfn2, and Opa1, while Fis1 and Dnm1l play key roles in mitochondrial fission. To investigate the expression of these key regulators during mitochondrial fission and fusion, we analyzed the expression levels of related proteins in the hearts of mice subjected to TAC-induced cardiac hypertrophy (Figure 3A).

**Figure 3.**
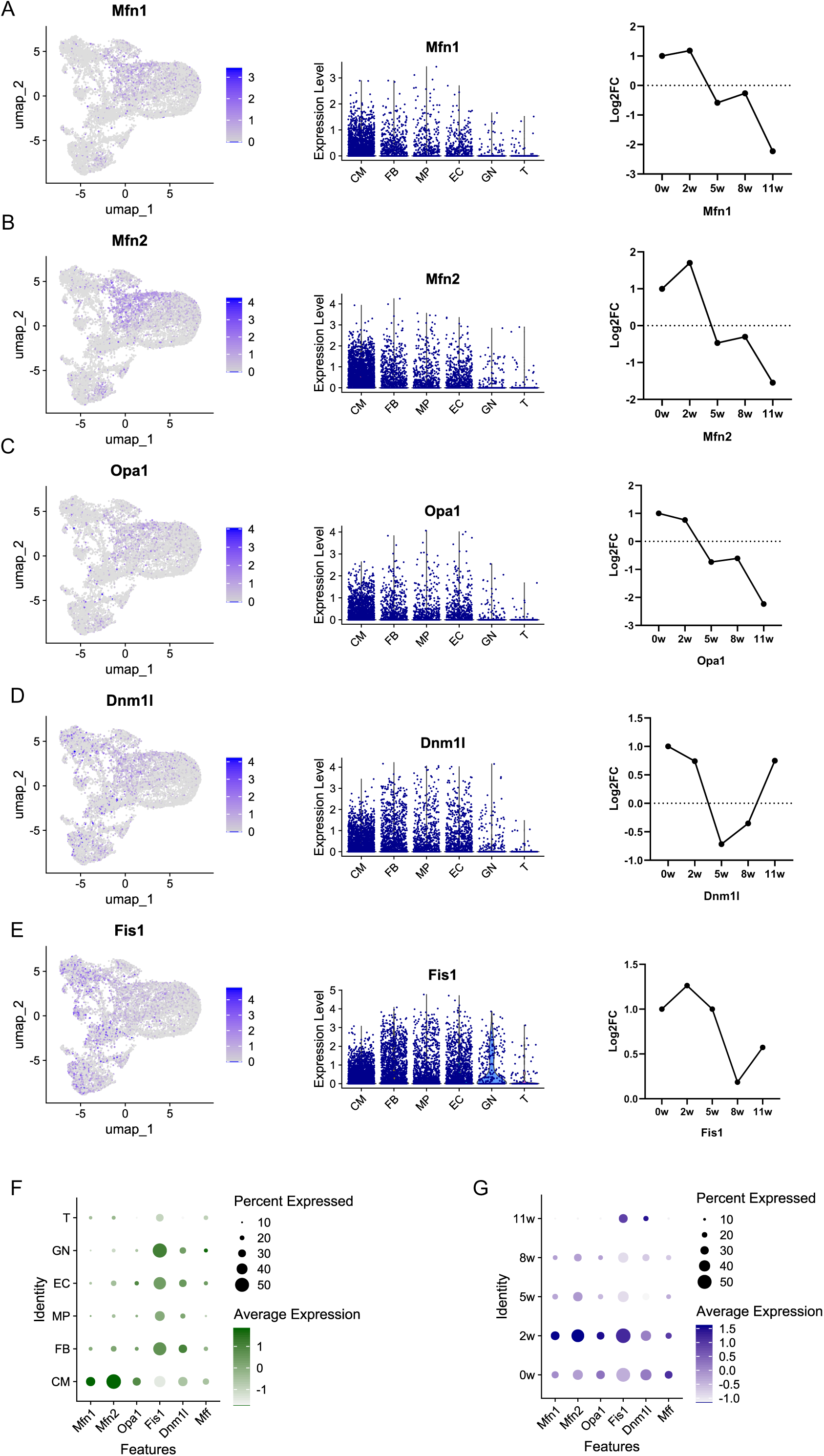
Expression of Mitochondrial Fusion and Fission Genes During Cardiac Hypertrophy. **A-E.** Left: UMAP plots showing the distribution of fusion and fission genes across cells. Middle: Gene expression profiles across different cell types. Right: Gene expression patterns at various time points. **F.** Dot plot illustrating the expression levels of mitochondrial fusion- and fission-related genes in different cell types. **G.** Dot plot displaying changes in the proportions of these genes over time, corresponding to different stages of disease progression. CM = cardiomyocyte; EC = endothelial cell; ECM = extracellular matrix; UMAP = uniform manifold approximation and projection; w = week.

Two weeks after hypertrophy onset, the expression levels of the fusion-related genes Mfn1, Mfn2, and Opa1 increased, but they gradually decreased during the later stages after TAC (Figure 3A, B, C, G).

In contrast, the fission-related gene Fis1 showed a consistent increase in expression throughout the postsurgery period. Dnm1l expression slightly decreased during the pathological hypertrophy stage but rose dramatically by week 11, coinciding with the transition to heart failure (Figure 3D, E, G). These findings suggest that mitochondrial fission plays a distinct role during the post-TAC stages.

In conclusion, both mitochondrial fusion and fission genes are preferentially expressed in myocytes. The expression of the fusion genes Mfn1, Mfn2, and Opa1 decreases during the pathological hypertrophy and heart failure stages, indicating a protective role. On the other hand, Dnm1l follows a biphasic pattern—decreasing initially but rising sharply during heart failure. Additionally, Fis1 expression increases throughout all time points, suggesting that while mitochondrial fusion decreases during pathological stages, fission tends to increase, reflecting a detrimental role.

### Mitochondrial Dynamics Enriched in Differentially Expressed Genes and Hub Modules During Physiological Hypertrophy

It is well established that regular exercise benefits cardiovascular health. Individuals who exercise regularly, including athletes, often exhibit physiological hypertrophy, a heart adaptation that is beneficial for overall health. In contrast, pathological hypertrophy—typically associated with conditions such as hypertension and heart failure—is detrimental. To explore the relationship between physiological and pathological hypertrophy, we analyzed bulk RNA-seq data from the hearts of exercise-induced model mice.

A total of 55,487 genes were selected for WGCNA. After the samples were clustered and the missing values were processed, an outlier sample was excluded. A soft-threshold power of b = 7 (scale-free R² = 0.85) was applied to ensure that the network followed a scale-free topology. Using a one-step method, we constructed a coexpression matrix, followed by dynamic hybrid cutting to identify 10 gene modules (Figure 4A). The distance matrix heatmap provided an overview of the similarities between samples (Figure 1B).

**Figure 4.**
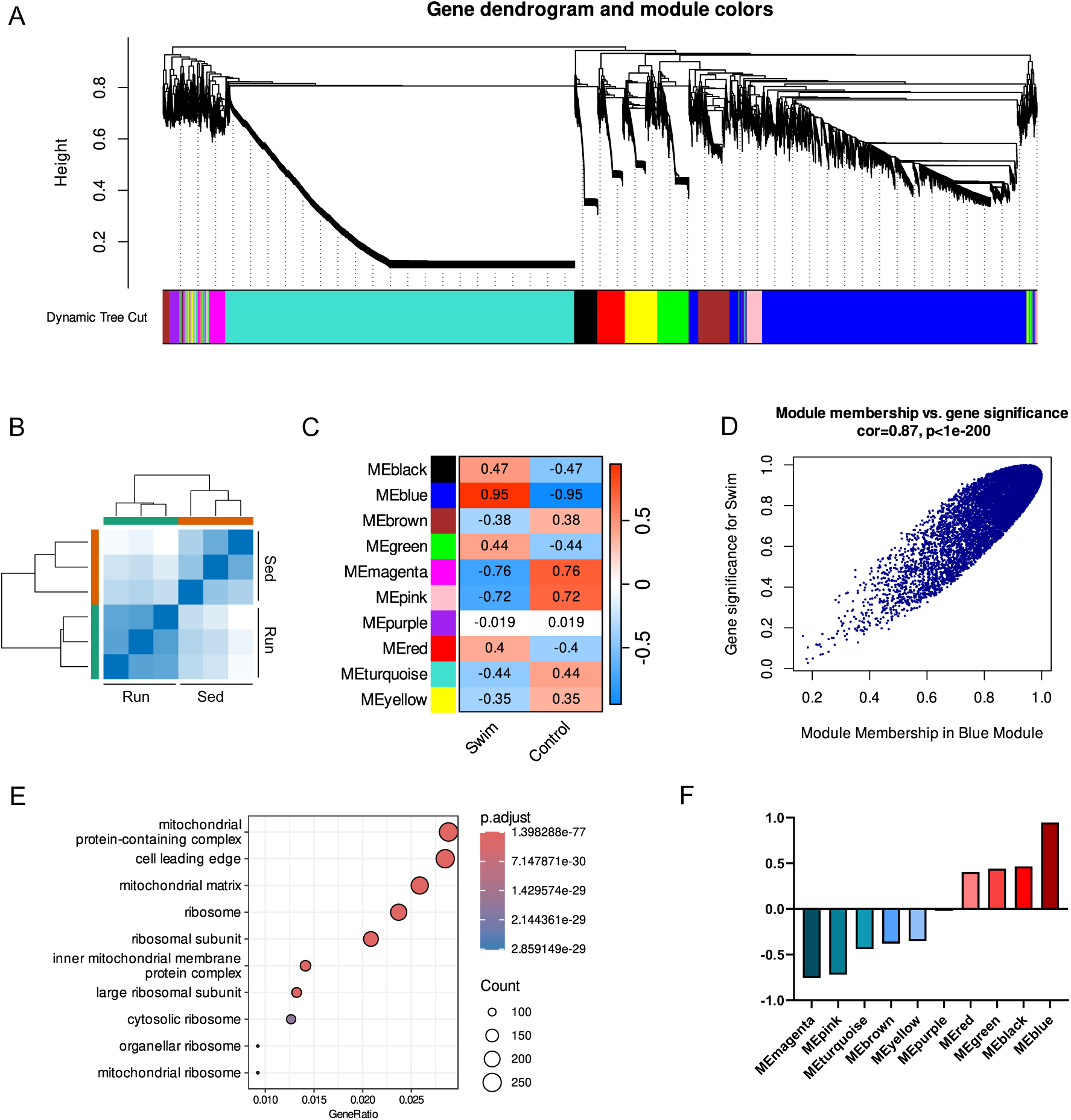
Co-Expression Network Construction and Hub Module Identification in the Exercise Mouse Model. **A.** WGCNA showing the hierarchical clustering dendrogram of coexpression modules, with each module represented by a distinct color. **B.** Correlation matrix illustrating the relationships between different samples. **C.** Heatmap displaying the correlation between hub genes and individual samples. **D.** Scatter plot highlighting the genes within the blue module associated with marker function. **E.** GO analysis of the blue module, highlighting enriched biological processes. **F.** Correlation scores of various coexpression modules; red indicates a positive correlation, whereas blue represents a negative correlation.

Among the identified modules, the blue module (comprising 248 genes) presented the strongest correlation with exercise conditions (Run group, cor = 0.95) (Figures 4C and 4F). Additionally, there was a strong correlation between gene significance (GS) and module membership (MM) within the blue module (cor = 0.87; P < e−200) (Figure 4D).

To further investigate the biological functions and signaling pathways associated with physiological hypertrophy, GO and KEGG pathway enrichment analyses were performed on the blue module genes. GO analysis revealed that the blue module was significantly enriched in mitochondrial-related components (e.g., the mitochondrial protein-containing complex, the mitochondrial matrix, and the inner mitochondrial membrane protein complex) and ribosomal structures (e.g., the ribosome subunit, the cytosolic ribosome, and the organellar ribosome) (Figure 4E).

In summary, WGCNA identified the blue module as the hub module most strongly associated with physiological hypertrophy. This module is enriched primarily in genes involved in mitochondrial and ribosomal functions, making it central to our subsequent analyses.

### Mitochondrial dynamics enriched in the DEG and Hub modules

A total of 4,733 differentially expressed genes (DEGs) were identified, including 2,036 upregulated and 2,697 downregulated genes, on the basis of adjusted P values < 0.05 and |logFC| > 0.5. These DEGs were visualized in a volcano plot (Figure 5A). Additionally, a subset of DEGs with the lowest P values is highlighted in a heatmap (Figure 5B).

**Figure 5.**
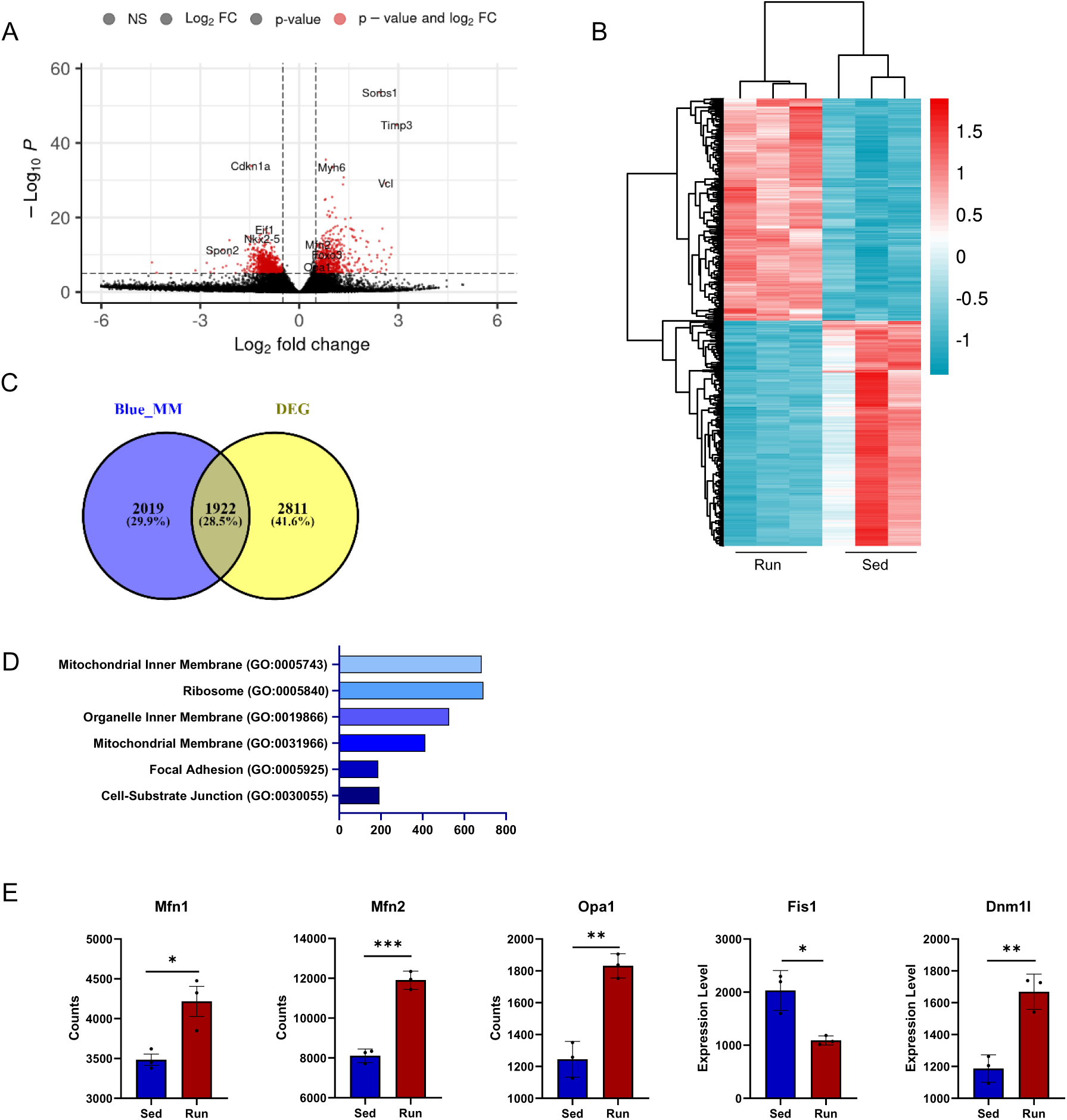
Mitochondrial Dynamics Enriched in Differentially Expressed Genes and Hub Modules During Physiological Hypertrophy. **A.** Volcano plot showing the results of differential gene expression analysis performed via the DESeq package. **B.** Heatmap displaying the significantly differentially expressed genes. **C.** Venn diagram illustrating the overlap between differentially expressed genes and genes in the blue module, with 1,922 overlapping genes. **D.** GO analysis of the 1,922 overlapping genes, highlighting enriched biological functions. **E.** Expression levels of fission- and fusion-related genes in exercise samples.

In the blue module, 3,767 genes with the highest connectivity were identified as candidate hub genes on the basis of screening criteria of |GS| > 0.80 and |MM| > 0.80. Among these, 3,264 genes overlapped with the DEGs, as shown in the Venn diagram (Figure 5C).

To better understand the functions of these intersecting genes, GO analysis was performed. The genes were significantly enriched in several cellular components, including the mitochondrial membrane (GO:0005743), ribosome (GO:0005840), organelle inner membrane (GO:0019866), focal adhesion (GO:0005925), and cell-substrate junction (GO:0030055) (Figure 5D).

Interestingly, blue module DEGs were enriched predominantly in mitochondria-related pathways, mirroring findings in the TAC model and highlighting the central role of mitochondria in cardiac physiology.

To further investigate mitochondrial dynamics, the expression levels of key genes involved in mitochondrial fusion and fission (Mfn1, Mfn2, Opa1, Fis1, and Dnm1l) were compared between the Run and Sed groups. The results revealed that Mfn1, Mfn2, Opa1, Fis1, and Dnm1l were upregulated, whereas Fis1 was downregulated in the Run group (Figure 5E).

In summary, analysis of the DEGs and hub modules associated with physiological hypertrophy revealed notable enrichment of mitochondrial dynamics pathways. These findings suggest that mitochondrial fusion is increased, whereas fission is decreased in exercise-induced cardiac adaptations.

### Validation of Mitochondrial Fission and Fusion Gene Expression in Exercise and TAC Models

To validate the findings from the previous analyses, we collected heart samples from both the TAC and exercise mouse models. The TAC model was studied 4 weeks postsurgery, whereas the exercise model involved mice starting swimming training at 8 weeks of age and swimming twice daily until they reached 12 weeks of age. After the heart samples were collected, RNA was extracted and reverse-transcribed into cDNA. PCR was then performed to measure the expression levels of the target genes, with 18S rRNA used as the control.

In the TAC model, Mfn1, Mfn2, Opa1, and Fis1 expression was significantly decreased, which was consistent with the single-cell RNA sequencing data from 2--5 weeks. Although Dnm1l exhibited a decreasing trend, the change was not statistically significant, suggesting a complex role for Dnm1l, which is involved in both mitochondrial fission and fusion. Notably, Dnm1l expression decreased at 5 and 8 weeks post TAC but dramatically increased by 11 weeks (Figure 6A).

**Figure 6.**
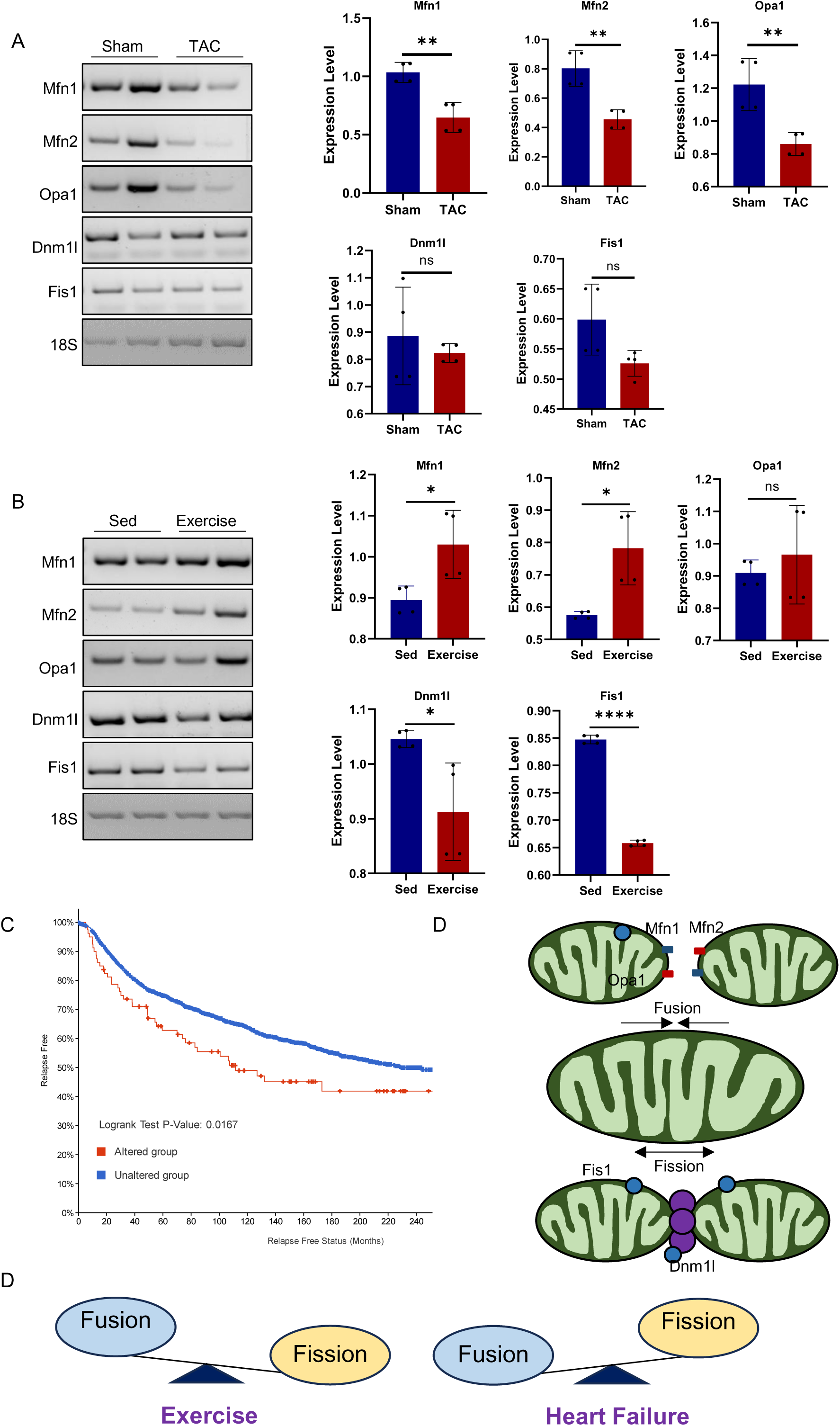
Validation of Mitochondrial Fission and Fusion Gene Expression in Exercise and TAC Models. **A.** Validation of mRNA expression levels in sham and TAC mouse hearts (n = 4 mice per group, *P < 0.05, **P < 0.01, Student’s t test). **B.** Validation of mRNA expression levels in sedentary and exercised mouse hearts (n = 4 mice per group, *P < 0.05, **P < 0.01, Student’s t test). **C.** Relapse-free survival curve for patients with Mfn1 mutation and cardiovascular disease. **D.** Schematic of mitochondrial fission and fusion dynamics. In exercise-induced (physiological) hypertrophy, fusion is increased relative to fission. In TAC-induced (pathological) hypertrophy, fission is increased relative to fusion.

In contrast, in the exercise model, Mfn1 and Mfn2 levels were significantly increased, which aligns with the results from the bulk RNA-seq data. While Opa1 showed an increasing trend, the change was not statistically significant, reflecting its diverse roles in both fission and fusion. Compared with those of TAC, Dnm1l and Fis1 were significantly downregulated, suggesting that mitochondrial fission has an opposing role during exercise (Figure 6B).

In women diagnosed with breast cancer, CVD is the leading cause of death, as cancer treatments can lead to early or delayed cardiotoxicity. We investigated clinical data on Mfn1 and its impact on relapse-free survival in breast cancer patients. As shown in Figure 6C, mutations in Mfn1 are associated with a shorter relapse-free period than mutations in the normal Mfn1 gene (Figure 6C).

Figure 6D illustrates the model of the mitochondrial fission and fusion processes. In the exercise model, mitochondrial fusion is upregulated, which likely enhances energy metabolism in myocytes. In contrast, mitochondrial fission is reduced, suggesting that fission may not benefit myocyte function under these conditions. Conversely, in the TAC model, fusion processes were reduced 4 weeks after TAC, although they increased at later stages.

In conclusion, mitochondrial fusion supports myocyte function and improves heart performance. Enhancing mitochondrial fusion may represent a potential therapeutic strategy for heart failure, as modeled in the TAC study.

## DISCUSSION

Understanding the mechanisms by which exercise promotes cardiac health, and whether these mechanisms can be effectively targeted, is of significant fundamental and clinical importance in cardiovascular diseases, including HF. The data presented here demonstrate that exercise dynamically regulates mitochondria-related pathways, which play pivotal roles in influencing HF outcomes, such as after TAC. Specifically, the mitochondrial fission and fusion pathways were upregulated at 2 weeks post TAC, but this activity decreased at 5, 8, and 11 weeks. Despite these trends in mitochondrial dynamics, fission-related genes behave differently under physiological and pathological conditions. Notably, in HFs, fission genes such as Fis1 and Dnm1l are upregulated, whereas fusion genes are downregulated, as expected. These findings suggest that Fis1 and Dnm1l play critical roles in the transition from physiological to pathological hypertrophy in cardiomyocytes.

Our findings underscore the importance of mitochondrial dynamics as a key modulator of cardiac growth and CM proliferation. These results indicate that mitochondrial dynamics, including fission and fusion processes, are increased during the early stages of cardiac hypertrophy but decrease later. This pattern can be interpreted as part of compensatory hypertrophy, akin to physiological hypertrophy. To our knowledge, these are the first studies to provide insight into mitochondrial dynamics from a single-cell perspective. Despite similarities in the expression patterns of fission and fusion genes between physiological and pathological hypertrophy, notable differences exist between the two conditions. The fission genes Fis1 and Dnm1l exhibited opposite expression changes, with upregulation in pathological hypertrophy. These findings suggest that Fis1 and Dnm1l are key drivers of pathological hypertrophy and heart failure, making them promising therapeutic targets for heart failure treatment.

Interestingly, prior studies lend experimental support to our findings, further validating the role of mitochondrial dynamics in cardiac pathology. Dnm1l is transiently activated 3 to 5 days after TAC but is chronically suppressed thereafter. The suppression of mitochondrial autophagy contributes to mitochondrial dysfunction and the progression of heart failure^18^. Additionally, the interaction between Dnm1l and Fis1 induces reactive oxygen species (ROS) production and excessive mitochondrial fragmentation, which can lead to cardiac disease^19^.

This work makes significant conceptual and practical contributions to the literature. First, through a comprehensive comparison of cardiac single-cell RNA transcriptomics across five time points, we identified a subset of pathways that were consistently altered across all stages. These pathways are enriched with functional candidates involved in pathological processes, suggesting promising therapeutic targets. Second, via WGCNA in exercise-induced model mice, we identified a blue module strongly correlated with swimming exercise. Enrichment analysis revealed that this module is associated with mitochondrial pathways and reflects the expression patterns observed in 2-week TAC models. Furthermore, we showed that the mitochondrial fission genes Fis1 and Dnm1l exhibit opposing expression patterns in physiological hypertrophy compared with heart failure, highlighting their potential as markers for distinguishing between adaptive and maladaptive cardiac remodeling.

Mitochondrial dynamics play a critical role in heart health and disease. Emerging research suggests that targeting these dynamics could be a therapeutic approach for various cardiac conditions, including myocardial infarction, hypertrophy, pulmonary arterial hypertension (PAH), ischemic heart disease, and HF. Although mitochondrial fusion is often considered protective and fission detrimental, this binary view may oversimplify their roles. Fission is essential for isolating damaged mitochondria for removal via mitophagy, and excessive inhibition of this process may worsen myocardial injury. Therefore, striking the right balance in regulating mitochondrial dynamics is essential for developing effective clinical applications.

## Conclusion

In summary, our studies provide novel insights into the critical role of the balance between mitochondrial fission and fusion in the transition from physiological to pathological heart hypertrophy. The fission genes Fis1 and Dnm1l exhibit opposing expression patterns in TAC-induced and exercise-induced cardiac hypertrophy, underscoring their roles as key drivers of heart failure. These findings suggest that Fis1 and Dnm1l could serve as potential diagnostic markers and therapeutic targets for managing heart failure.

## Abbreviations

HF: Heart failure
TAC: Transverse aortic constriction
CVD: Cardiovascular disease
scRNA-seq: Single-cell RNA sequencing
CMs: Cardiomyocytes
Ecs: Endothelial cells
FBs: Fibroblasts
MPs: Macrophages
GNs: Granulocytes
Mfn1: Mitofusin 1
Mfn2: Mitofusin 2
Opa1: Opticatrophy protein 1
Fis1: Mitochondrial fission 1
Dnm1l: Dynamin 1-like
EF: Ejection fraction
MM: Module membership
DEGs: Differentially expressed genes
GS: Gene significance

## Declarations

## Ethics approval and consent to participate

All animal procedures were performed in accordance with protocols approved by the Institutional Animal Care and Use Committee (IACUC) of Shanghai University, China.

## Consent for publication

Not applicable.

## Availability of data and material

All data associated with this study are present in the paper or in the Supporting Information. The scRNA-seq data of keloids from the published paper are available in the GEO under accessions GSE120064 and CRA007207. Data, codes, and materials will be made available upon request.

## Author contributions

Gao led the data analysis and manuscript writing. All the authors reviewed and approved the final version of the manuscript.

## Acknowledgments

This manuscript was supported by Dr. Xiao, Dr. Li and Dr. Gao.

## Authors’ information

^1^Cardiac Regeneration and Aging Lab, Institute of Geriatrics (Shanghai University), Affiliated Nantong Hospital of Shanghai University (The Sixth People’s Hospital of Nantong) and School of Life Science, Shanghai University, Nantong 226011, China. ^2^Institute of Cardiovascular Sciences, Shanghai Engineering Research Center of Organ Repair, Joint International Research Laboratory of Biomaterials and Biotechnology in Organ Repair (Ministry of Education), School of Life Science, Shanghai University, Shanghai 200444, China. ^3^Shanghai University Hospital, Shanghai University, Shanghai 200444, China. ^4^Department of Internal Emergency Medicine, Shanghai East Hospital, School of Medicine, Tongji University, Shanghai, 200120, China. ^5^Department of Cardiology, Tenth People’s Hospital, School of Medicine, Tongji University, Shanghai, 200090, China.

